# Bacttle: a microbiology educational board game for lay public and schools

**DOI:** 10.1101/2024.04.04.588042

**Authors:** Tania Miguel Trabajo, Eavan Dorcey, Jan Roelof van der Meer

## Abstract

Inspired by the positive impact of serious games on science understanding and motivated by personal interests in scientific outreach, we developed ‘Bacttle’, an easy-to-play microbiology board game with adaptive difficulty, targeting any player from 7 years old onward. Bacttle addresses both the lay public and teachers for use in classrooms as a way of introducing microbiology concepts.

The layout of the game and its mechanism are the result of multiple rounds of trial, feedback and re-design. The final version consists of a deck of cards, a 3D-printed board and tokens (with a paper-based alternative), with all digital content open source. Players in Bacttle take on the character of a bacterial species. The aim for each species is to proliferate under the environmental conditions of the board and the interactions with the board and with other players, which vary as the play evolves. Players start with a given number of lives that will increase or decrease based on the traits they play for different environmental scenarios. Such bacterial traits come in the form of cards that can be deployed strategically.

In order to assess the impact of the game on microbiological knowledge, we scored differences in the understanding of general concepts before and after playing the game. We assessed a total of 169 visitors at two different university open day science fairs. Players were asked to fill a brief survey before and after the game with questions targeting conceptual advances. Results show that Bacttle increases general microbiology knowledge on players as young as 5 years old, and with the highest impact on those who have no *a priori* microbiology comprehension.

## 2. Introduction

The first written definition of serious games was given by Clark Abt back in the seventies, who proposed board and card games as a tool to improve education and prevent academic failure among students. He defined them as games carefully designed to not just entertain, but with a clear educational purpose(1). Nowadays, the term ‘serious games’ refers to video games that fulfil such educational purposes, developed to play virtually through a screen(2,3). They feature a goal to be reached, constrained by rules and limitations on what a player can do, with a sense of competition while maintaining a playful aspect. Such aspects are part of the original definition of serious games and thus, despite most of current research being focused on digital games, the same considerations apply to board and card games.

According to a meta-analysis carried out by Riopel(4), science-related serious games improve three different cognitive learning outcomes when compared to more conventional instructional methods, namely (i) the declarative knowledge (reflected by post-tests answered immediately after playing the game), (ii) knowledge retention (delayed post-tests) and (iii) the application of such knowledge when performing a task. Many of the articles reviewed assert that such practices benefit from the additional support of other educational tools, such as teachers’ guides, theoretical content, and the class environment(5,6). These results are consistent with the review of Rutten(7), where serious games were analysed as an add-on in traditional education, showing improvement of learning outcomes, conceptual understanding, and predictive ability.

Learning outcomes of serious games do not depend on the scientific area they cover (e.g., engineering, biology, physics)(4). A variety of content has been developed in the area of microbiology serious games, but mostly, their educational effects have not been measured and thus we cannot conclude on their impact on learning. Microbiology-based serious games revolve around the relevance of microbes for human health and often highlight their negative aspects. For instance, the board game MyKrobs (created by Gilbert Greub; https://mykrobs.ch/en/) educates players on multiple microbe species that endanger human health. It raises awareness about the diseases they cause and how to prevent their transmission. Its targeted audience ranges from young adults to adults. Another example is Gut Check, created by Daniel Coil, where the goal is to build a healthy gut microbiome while disrupting those of the player’s opponents(8). This game makes an effort to put into perspective the microbial world in terms of “good” and “bad” microbes, showcasing the fact that not all of them are harmful. The game Gutsy, which was released by the American Museum of Natural History (https://www.amnh.org/explore/ology/microbiology/gutsy-the-gut-microbiome-card-game) follows the same topic of the gut microbiome, again with the focus on humans. Lastly, the board game Strain, published by HungryRobot, features the concept of resource management, attack and defence, while putting together the individual player’s perfect organism (the game has been discontinued, no reference exists). Some other microbiology-based serious games exist that were specifically created for classroom usage, for instance, Outbreak!(9), Biofilm building(10) or MedMyst(11). Of all the games cited above, only MyKrobs is currently available for purchasing, while the rest has been discontinued or is available as open source material that can be printed at home.

In the present work, we propose an easy-to-play serious board game named ‘Bacttle’ that showcases the life of microbes in the environment and targets players in primary school. The game covers basic concepts of environmental microbiology, starting from the concept that microbes are found all around us (i.e., soils, rivers), and have very different properties, to more specific biological information, such as motility or biofilm formation. We wanted the game to illustrate the many challenges bacterial species face in their survival and proliferation, from being threatened by viruses (bacteriophages) to undergoing constant competition for resources with other species. Although our main target is primary school kids, thanks to the game’s adaptive difficulty, both young (starting from age 7) and older players can learn about microbiology facts, while enjoying the game’s strategic challenges.

As suggested by Wouters(12), serious games show even better results when played in groups rather than individually. The prototype version of Bacttle is intended for up to six players. Likewise, following the conclusions from the meta-analysis of Riopel(4) that there is no correlation between how realistic a game interface is and how much is learned from it, Bacttle cards and board are illustrated with a cartoonesque touch. This makes it easier to depict the meaning of the traits in the cards using the least amount of text, whereas humanizing some aspects of the scientific definitions serves to facilitate concept transmission to the players.

Our initial hypothesis was that playing Bacttle would increase general knowledge on bacterial life among players irrespective of their age. To evaluate this, we measured the declarative knowledge of players with varied background (i.e., age, base knowledge) before and after playing the game.

Bacttle will be made publicly available as both an open-source downloadable version and as a purchasable physical version. We hope that the game will entertain players of a wide age range through its playful engagement of microbiology concepts, and secondly, that it can support microbiology teaching at schools.

## 3. Materials and methods

### 3.1. Game elements and gameplay

The original motivation behind Bacttle was to offer an educational activity within the *Mystères de l’UNIL* public exhibition in 2022, taking place at the University of Lausanne. This open and free annual event showcases different activities throughout the campus, where all researchers can share their research with the lay public. Due to the large positive feedback of the workshop, we developed the idea into a game that would transmit the same original notions about microbiology but with a self-sustained gameplay. Over the course of six months, the game was developed and refined. The

first versions of the game were printed on paper (board and tokens included). This allowed the gameplay to be improved, new cards to be created and the spatial strategy to be added as an advanced rule for the game. During a second phase of testing, the 3D printed copies of the board and tokens were designed and different trial versions were printed to check for the optimal size and features, while using biodegradable polymer of different colours.

The final version of the game includes a deck with 34 different cards: 17 environmental scenarios to play against (sized 79x120 mm) + 17 traits that can be strategically played by each player (63x88 mm). Each trait is represented four times, resulting in a deck with 85 cards (Fig. 1).

**Figure 1.**
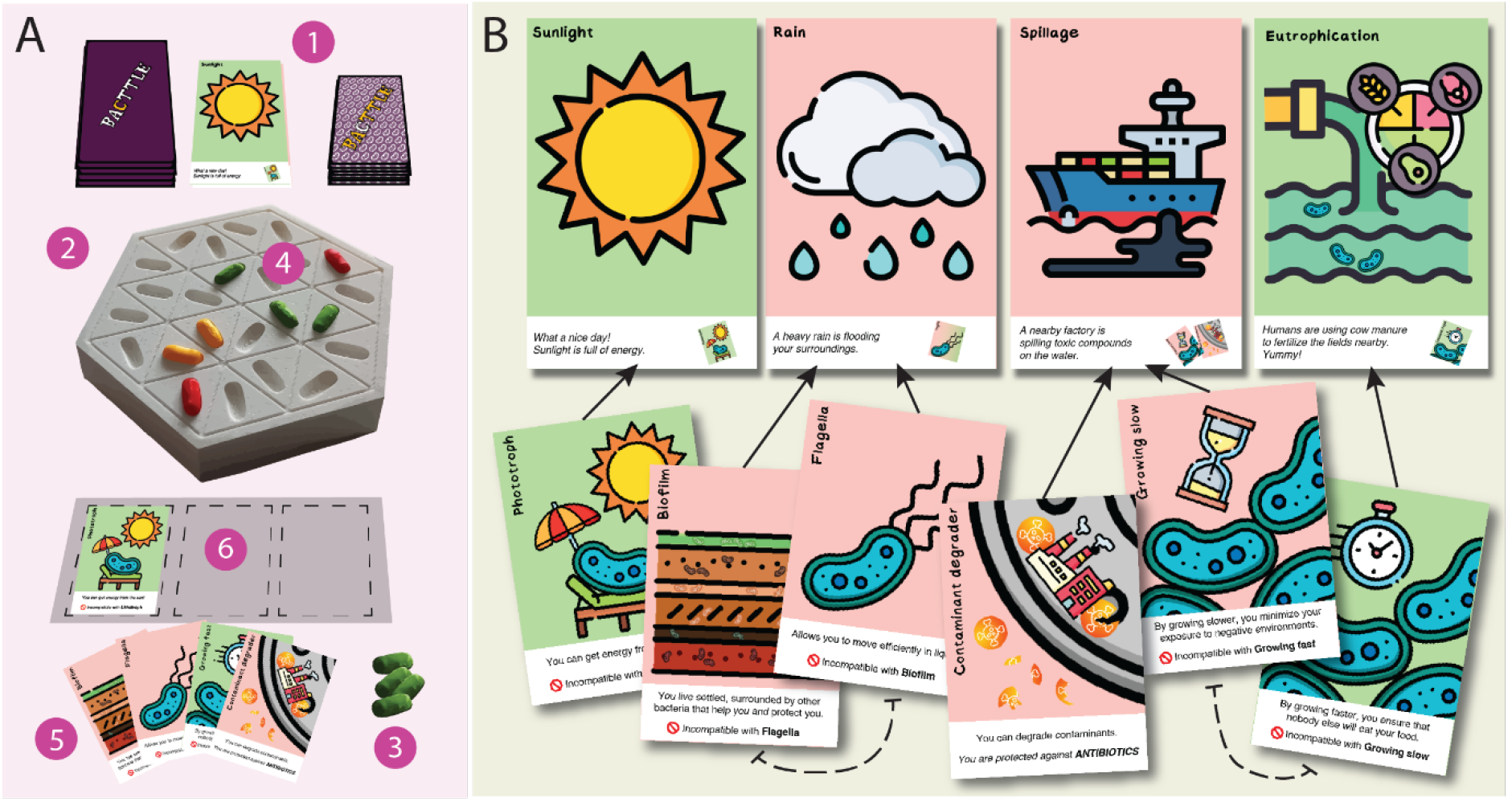
Overview of Bacttle. A. Set up of the game: 1) Card deck with the challenges to be unraveled and card deck with the traits. 2) Board support for the tokens, representing the habitat where the player’s microbes are living. 3) Tokens representing the different bacterial species for each of the players. 4) Every player starts with three tokens (cells) on the board, which can increase or decrease during the game’s rounds. 5) Players always have four trait cards in their hands. 6) Players can play three trait cards in front of them. B. Example of environmental challenges (top) and trait cards (bottom). Green environmental challenges represent benign situations that can be beneficial with the appropriate trait card (multiplying tokens), while red represent harmful situations that require a trait in order to preserve the tokens in play. Arrows relate to the trait cards that are useful for every environmental challenge. Some trait cards cannot be played simultaneously (black dotted lines), highlighted by their red inhibition symbol. Credit: illustrations were made using freely available icons from Freepik, ultimatearm, kmgdesign, and Smashicons.

In Bacttle, each player is a bacterial species that lives in the environment (the board support, Fig. 1A). Each player (species) starts with the same number of cells (the bacterial tokens). The number of cells per species on the board represents the number of lives the player has in the game. The goal is to be the player with the highest number of cells in play by the end of the game. Every round of the game develops around a new challenge, played in form of cards (Fig. 1B). Some challenges are detrimental (red cards) and will decrease the number of cells, unless the player can protect them by using trait cards (Fig. 1B). Other challenges give the opportunity to increase the number of cells (positive challenges, green). Lastly, battle challenges allow players to attack each other using their biological ‘weapons’ (yellow cards, not shown). The game ends once all the challenges have been played or only one species remains on the board.

The challenges illustrate habitat conditions for microbial life, induced either by natural or human causes. For instance, ‘Sunlight’ is a positive natural challenge for the habitat, which allows only phototrophic species to multiply. In contrast, ‘Rain’ is a potential negative natural challenge, which will wash away those species from the board that cannot swim or form a sticky biofilm (Fig. 1B). As examples of human-induced challenges, ‘Eutrophication’ allows fast growing species to multiply thanks to the added high nutrient levels, while ‘Spillage’ intoxicates all species unless they can degrade the contaminants or slow down their growth (Fig. 1B).

The abilities of species to photosynthesize or to swim, to form a biofilm or to degrade contaminants, come in the form of trait cards that players draw and have in their hand. The complexity of the game resides in the fact that players can only use three trait cards for their species at a time. Some trait cards are useful for multiple environments or challenges, but some trait combinations are incompatible (Fig. 1B). Each round reveals a new challenge (environmental card, Table 1), and players have the chance to play a trait card (Table 2) from their hand to be used in their favor. Some traits are special and allow players to carry out actions and interact with other species. For instance, if a player plays the card ‘Sporulation’, the species is protected against any negative challenge during that round. The card ‘Mutation’ allows one player to sabotage another player by removing a key trait from the other player’s species. Lastly, some trait cards can be played to equip the player’s species with biological ‘weapons’, which can be used to diminish the lives of competitor species. As examples, by the release of antibiotics or of bacteriocins, or by using a type VI secretion system (stabbing other cells, Table 2).

**Table 1.** Environmental cards and their description.

| *Environmental card* | *Effect* | *Description* |
| --- | --- | --- |
| *Rain* | - | Heavy rain changes the structure of the top soil, which can be detrimental specifically when there is a lack of vegetation. Heavy rain also leads to flooding. |
| *Drought* | - | Drought impacts microbial life by the risk of desiccation. |
| *Pollution* | - | Increase of harmful compounds in the air, which can be toxic for microorganisms. |
| *Spillage* | - | Increase of harmful compounds in the water, which can be toxic for microorganisms. |
| *Viral attack* | - | Threat of infection by bacteriophages, which can kill the cell. |
| *Food deprivation* | - | Lack of nutrients; leading to stalling of growth and starvation. |
| *Predation* | - | Killing of microorganisms by swallowing and ingestion of the cell through larger eukaryotic unicellular organisms or nematodes that feed on bacteria. Can also occur as a consequence of attack by other bacteria, which leads to lysis of the cell and feeding on its content. |
| *Fertilization* | + | The addition of fertilizer to the soil by humans. This leads to an overabundance of certain nutrients (notably phosphate and ammonium), which are otherwise limiting proliferation of certain groups of microorganisms. |
| *Sunlight* | + | Use of light as energy for growth. Can allow certain microorganisms to proliferate on abundant carbon dioxide, which otherwise does not contain sufficient energy to sustain building new cells. This process is known as photoautotrophic growth. |
| *Eutrophication* | + | Refers to a state of overabundance of nutrients in the environment (see Fertilization), which can lead to excessive growth of photoautotrophic microorganisms (e.g., algae, cyanobacteria, dinoflagellates). The secondary effect of this can be formation of toxic compounds (e.g., released by cyanobacteria) or loss of oxygen. |
| *Cross feeding* | + | When living together in a biofilm, bacteria share resources like in a food chain, where the by-products of one species metabolism can be still useful for another species, and so on. |
| *Minerals* | + | Use of chemical energy liberated from the conversion of inorganic minerals as power source for growth. Can also allow certain microorganisms to proliferate on carbon dioxide. |
| *Rhizosphere* | + | Plants sustain microbial communities by exudation of food and nutrients through the roots. Microorganisms living close to the plant root (the rhizosphere) can profit from this supply. |

**Table 2.** Trait cards and their targeted notions.

| *Trait cards* | *Useful against or in combination with* | *Targeted notion* |
| --- | --- | --- |
| *Flagella* | Rain, Predation, Food deprivation | Bacteria become motile with flagella. This allows them to move through their environment, and find specific nutrients, or move away from toxic compounds. Works specially in aqueous environments or when small water films cover particles in soil. |
| *Biofilm* | Cross feeding | Having the trait to form Biofilms, allows the bacteria to profit from the Flow environment card. |
| *Phototroph* | Sunlight | Having this trait in combination with Sunlight as environment, allows Phototrophs to utilize the light as energy source. |
| *Lithotroph* | Minerals | Having this trait in combination with Minerals as environment, allows Lithotrophs to use minerals as energy source. |
| *Viral immunity* | Viral attack | Having the Immunity trait card allows bacteria to prevent or defend against phage attack. |
| *Toxin immunity* | Bacteriocins, Antibiotics | Having this card allows protection against harmful compounds produced by others. |
| *Capsule* | Drought, Viral attack | Being able to produce a capsule helps bacteria to protect from Drought conditions, because the cell will not lose its water content. Capsule also prevents attack by phages. |
| *Growing slow* | Drought, Pollution, Spillage, Food deprivation | Slow growth helps bacteria to save energy and resources. This can make them more resilient to harmful compounds and harsh conditions. |
| *Growing fast* | Fertilization, Eutrophication, Rhizosphere | Faster growth gives advantage to certain bacterial species which can thrive in conditions of abundant nutrients. This allows them to proliferate quickly and colonize the habitat. |
| *Contaminant degrader* | Pollution, Spillage | Having this card allows the bacteria to take up harmful compounds and break them down. This gives them energy for growth and leads to detoxification of the environment. |
| *Bacteriocins* | Other players | This trait allows the bacteria to produce bacteriocins, which are substances that kill or disrupt the growth of other bacteria. Bacteriocins can be very specific against certain groups of other bacteria. |
| *Piercing weapon* | Other players | This trait card allows the bacteria to build injection needles, which can be used to pierce through the cell envelope of other bacteria and inject toxins, or directly disrupt the cell. These needles are known as type VI secretion systems. |
| *Antibiotics* | Other players | Allows the bacteria to produce antibiotics: powerful compounds that inhibit the growth of other bacteria or kill them entirely. Antibiotics frequently have a wide spectrum of action and can target many groups simultaneously. Bacteria can acquire resistance to antibiotic action by various means. |
| *Conjugation* | - | This card allows bacteria to share traits with others. It refers to a process by which cells from different species transmit part of their genes almost through a tube that temporarily connects them. |
| *Transformation* | - | This card allows bacteria to implement lose traits as their own. It refers to a process that allows some bacteria to take up DNA from the environment and implement it into their own genetic material by recombination. |
| *Mutation* | - | This card will deactivate another trait. It refers to a spontaneous process by which genetic information in the cell slightly changes, which can affect the functions encoded by genes. For instance, a mutation could lead to a different amino acid being incorporated in a protein, or to shortening of the protein. |
| *Sporulation* | - | This card allows the bacteria to form spores. By doing so, they become metabolically dormant but highly resistant to harsh conditions. Spores cannot multiply. |

### 3.2. Educational aspect

The environments displayed in this first edition of Bacttle are limited to habitats and situations, which are impacted by human activity. Table 1 summarizes the environmental cards and their meaning.

The traits that are showcased in Bacttle consist of a mixture of physiological and structural characteristics that the species can obtain in order to enhance their performance. The characteristics introduce microbiology-related terms in a playful manner without becoming too technical. For instance, the type VI secretion system(13) is called ‘piercing weapon’ to ease the understanding of what this trait implies. Table 2 summarizes the trait cards and the notion we aimed to target. The notions to be learnt through playing come from both the text displayed on the cards and the illustrations themselves.

### 3.3. Participants in the study

Prototypes of the game were tested and evaluated on two different occasions: (i) the Mystères de l’UNIL 2023 (science fair organised by the University of Lausanne -UNIL, in total four days), and (ii) Scientifica 2023 (a science fair organised by the Swiss Federal Institute of Science and Technology, ETH Zurich; one day). Prototypes were language-adapted to the public; for Mysteres de l’UNIL the cards were in French, and for the Scientifica public the cards were translated into German.

The first two days at Mystères de l’UNIL were reserved to organised school class visits, under direction of the class teacher and with a strict time limit. Schoolkids played the game in groups of four and had a window of 15 minutes to learn how to play and try the game. The range of ages varied from 8 to 13 years old (primary school). The last two days at Mystères de l’UNIL and the day at Scientifica were open to general lay public. In these cases, groups were formed spontaneously, leading to parents often playing alongside their children. For both groups (schoolkids and lay public) the same game restrictions were applied for consistency of the data: preparation and explanation of the rules, filling the first survey, playing the game for 15 minutes, and filling the second survey. For the lay public, the range of players’ ages was 4 to 53 years old. It was observed that in a limited number of cases, when families were playing together, parents would help answer the evaluation forms of the young kids. We tested for such bias by treating the surveys of schoolkids separately from those of lay public.

Data from both events were grouped (Lausanne and Zurich). In total, 64 surveys were returned in the category ‘schoolkids’, and 88 surveys in the category ‘lay public’.

### 3.4. The survey

In order to assess the declarative knowledge gained by playing the game, a brief evaluation survey was designed to be answered anonymously before and after playing the game. The survey (Table 3) consisted of 16 different questions to gather information about the players’ age and their initial level of knowledge in microbiology, acquired knowledge from the game, and appreciation of the game.

**Table 3.** Evaluation survey content.

| *ID<sup>1</sup>* | *Question text* | *Possible answers<sup>2</sup>* |
| --- | --- | --- |
| *A* | How old are you? |  |
| *B* | Do you know what a bacterium is? | y/n |
| *C* | Do you know what a bacterial capsule is? | y/n |
| *D* | Do bacteria have tools to harm each other? | y/n/idk |
| *E* | Do bacteria reproduce at the same pace? | y/n/idk |
| *F* | What is sporulation? | A resistant state that some bacteria can achieve under unfavorable conditions.<br>The release of toxins by bacteria.<br>idk |
| *G* | What are flagella used for? | Sticking to surfaces.<br>Motility in liquid environments.<br>idk |
| *H* | What does it mean to be lithotrophic? | A bacterium can get energy from minerals.<br>A bacterium can get energy from the sunlight.<br>idk |
| *I* | Can bacteria be infected by viruses? | y/n/idk |
| *J* | Are all bacteria harmful for humans? | y/n/idk |
| *K* | How many bacteria are in a coffee spoon of yoghurt? | Millions<br>Hundreds<br>idk |
| *L* | How easy did you find the gameplay? | VE/E/A/D/VD |
| *M* | Did you find the card content easy to understand? | VE/E/A/D/VD |
| *N* | Did you like the setup of the game? | y/n/idk |
| *O* | Would you like to play this game again? | y/n/idk |
| *P* | What can we improve? |  |

1) Question A categorizes the player's age; B and C assess the initial level of knowledge in microbiology (none -both questions are answered negatively-, basic -player knows what a bacterium is but not a bacterial capsule-, or advanced - both answers are positive-); questions D-I score knowledge acquisition; J and K are control questions; L-O evaluate the appreciation of the game; and P is an optional free text-entry answer for additional feedback.
2) y= yes, n=no, idk=I don't know, VE=very easy, E=easy, A=adequate, D=difficult, VD=very difficult.

### 3.5. Statistical analysis

A *posteriori* data analysis was carried out with R version 4.1.0 through RStudio version 1.4.1106. The improvement score was calculated by counting +1 point for a change of a wrong or ‘I don’t know answer’ (in the pregame questionnaire) to a correct answer; –1 for a correct or ‘I don’t know answer’ changing into an incorrect reply after the game; and a 0 for any other combination of answers.

Incomplete surveys were discarded (13% of the total).

Differences in the improvement score between the groups of schoolkids and lay public were tested by a Wilcoxon rank sum test. Knowledge acquisition was tested using a Wilcoxon signed-rank test, by using as a control for comparison the score from questions unrelated to the content of the game. Control questions were microbiology-related but their answer could not be learnt from playing the game.

To study the relation between the score and social variables like age or initial knowledge on the subject before playing the game, two additional analyses were carried out. The correlation between age and improvement scores was examined using Pearson’s linear correlation, while differences in the distributions of scores and initial knowledge were tested using a Kolmogorow-Smirnow test.

### 3.6 Simulation

In order to verify that results were not likely to be the consequence of random answering, we simulated a set of data with the same size as the number of returned surveys, where answers to the knowledge acquisition questions and the control questions (questions D-K, Table 1) were randomly answered. A Wilcoxon signed-rank test was then used to compare the knowledge acquisition with the control data.

The simulation assumed a discrete uniform distribution between right/wrong/’I don’t know’ answers and considered each question to be independent of each other (identically independently distributed).

## 4. Results

### 4.1. Learning outcomes

We found a significant increase in the improvement score for the test questions compared to the control questions after playing the game (Fig. 2, p-value=2.5x10^-12^ for the schoolkids category and p-value=5.8x10^-8^ for the lay public category). These results imply that Bacttle improves the declarative knowledge of players.

**Figure 2.**
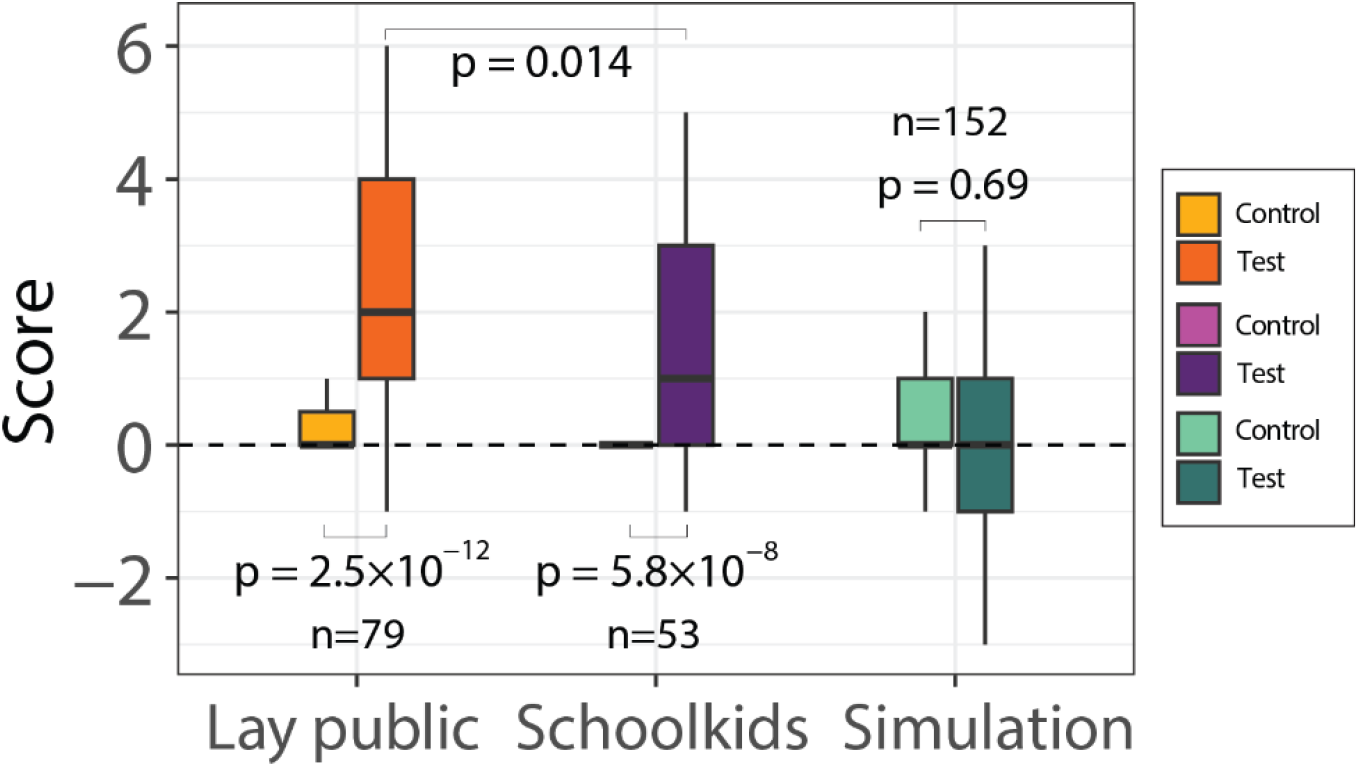
Playing Bacttle improves self-declared microbiology knowledge. Box plots show the score ranges of control questions (left bars) and knowledge acquisition questions (right bars) for the two different surveyed groups and a random reply simulation. Light colors of pairs point to control questions. n=number of included surveys. P-values between control and sample stem from a paired Wilcoxon signed rank test, while p-values between lay public and schoolkid samples result from a Wilcoxon rank sum test. Box plots show the lower and upper quartiles and the median, plus the outlier range (5^th^ to 95^th^ percentiles).

Lay public scored significantly higher than schoolkids (Fig. 2, p-value=0.014), confirming a potential bias among both groups. Such difference may be due to the fact that parents were supporting their children by reading out loud the questions and even providing feedback in some cases, increasing the chances that an *a priori* wrong or ‘I don’t know’ answer would become an *a posteriori* correct answer. The difference could also be the consequence of having a wider range of ages in this group, which implies more adults answering the surveys, who are likely to pay more attention to the game content and questions and obtain a higher score.

The simulated data scored on average a zero (Fig. 2), meaning that there was no increase nor decrease in knowledge by random answering. Based on the random simulation, 26% of the players would have been expected to have an improvement score ≥1. In the actual survey, we observed that 77% of the participants scored ≥1, supporting the learning effect. Lastly, the scores on control and knowledge acquisition questions were not significantly different for the random replies, as expected (Fig. 2, p-value=0.69). We can thus conclude that the game indeed objectively improved self-declared microbiology knowledge.

### 4.2. Correlations with age and previous microbiology knowledge

In order to corroborate if the player’s age affects the learning outcome, we analysed the correlation between age and improvement score (Fig. 3A). The range of players’ ages varied from 4 to 53 years, with a high prevalence of 10-year-old players, which was the most common age among the schoolkids. We found no significant correlation between age and score (Pearson correlation coefficient = -0.077, p-value=0.38), suggesting that the game’s educational aspect does not depend on the age of the player.

**Figure 3.**
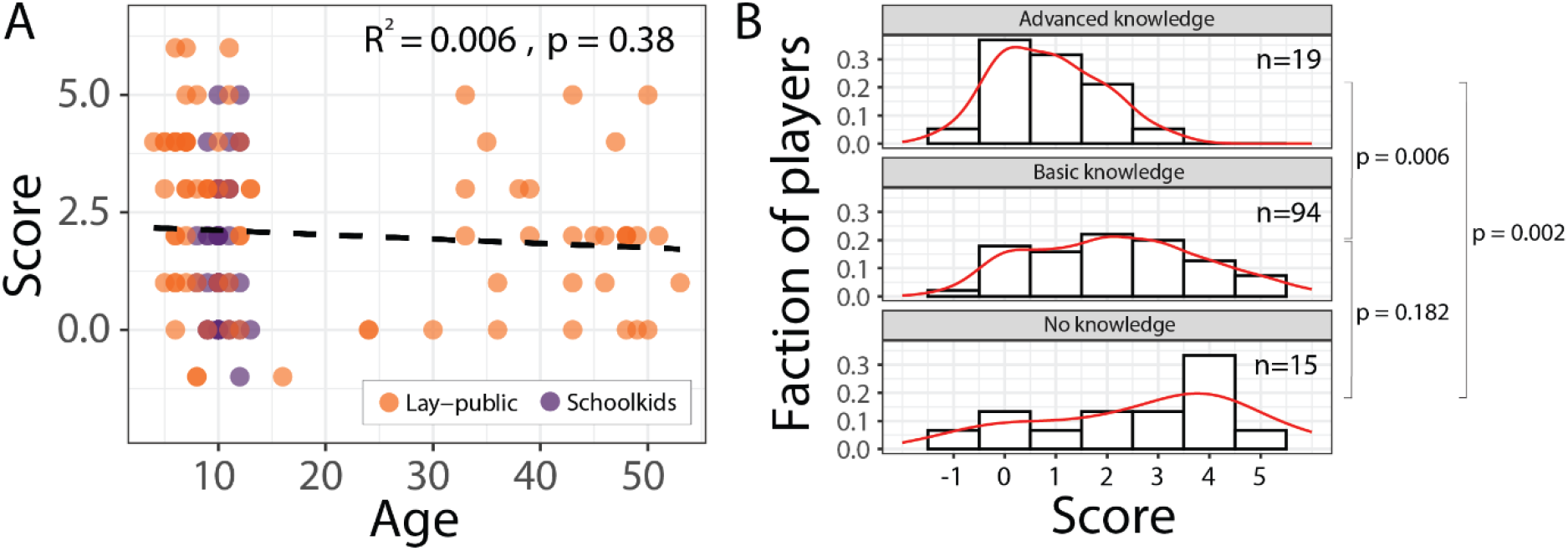
Analysis of the effect of age or basic microbiology knowledge on the improvement score. A. Pearson correlation between the score and the player’s age (R-square=-0.077, p-value=0.38). B. Score distributions and comparison by knowledge categories using Kolmogorow-Smirnow testing. ‘no knowledge’ vs. ‘basic knowledge’; p-value=0.182. ‘basic knowledge’ vs. ‘advanced knowledge’; p-value=0.006. ‘no knowledge’ vs. ‘advanced knowledge’; p-value=0.002).

Finally, we analysed the potential correlation between improvement score and *a priori* microbiology knowledge, under the assumption that players with a higher initial knowledge in microbiology will learn less (lower score) than those with only basic or no previous knowledge in microbiology (higher score). This categorization is made based on two yes-or-no questions (Table 3, questions B and C).

Players replying negatively to question B, independently of their answer to question C, are considered as falling in the category ‘no previous knowledge’. If they answered ‘yes’ to question B and ‘no’ to question C, they are categorized as having ‘basic knowledge’. Lastly, those players answering positively both questions are considered with ‘advanced knowledge’.

Indeed, when plotting the improvement score distributions as a function of declared initial knowledge, a tendency can be seen (Fig. 3B). For players declaring no previous knowledge in microbiology, the abundance of improvement scores peaks at four. On the opposite side, players with declared advanced knowledge have the highest prevalence of a zero-improvement score.

Players with declared basic knowledge show a homogeneous distribution of improvement scores with an average score of two. Such differences on the distributions were confirmed by the Kolmogorow-Smirnow test, which showed a significant difference between the distributions of ‘basic knowledge’ and ‘advanced knowledge’ (p-value=0.006) and ‘no knowledge’ and ‘advanced knowledge’ (p-value=0.002). However, no significant difference was found between the groups ‘no knowledge’ and ‘basic knowledge’ (p-value=0.182). This may be due to the fact that the sample size is too small to make accurate comparisons.

### 4.3. Appreciation of the game

Most players qualified the game’s difficulty as ‘Adequate’, followed by ‘Easy’ (Fig. 4A). A similar trend was observed to describe the level of the readability and the ease to understand the text on the cards (Fig. 4B). Almost all players (94%) stated that they liked the game and 86% would play it again.

**Figure 4.**
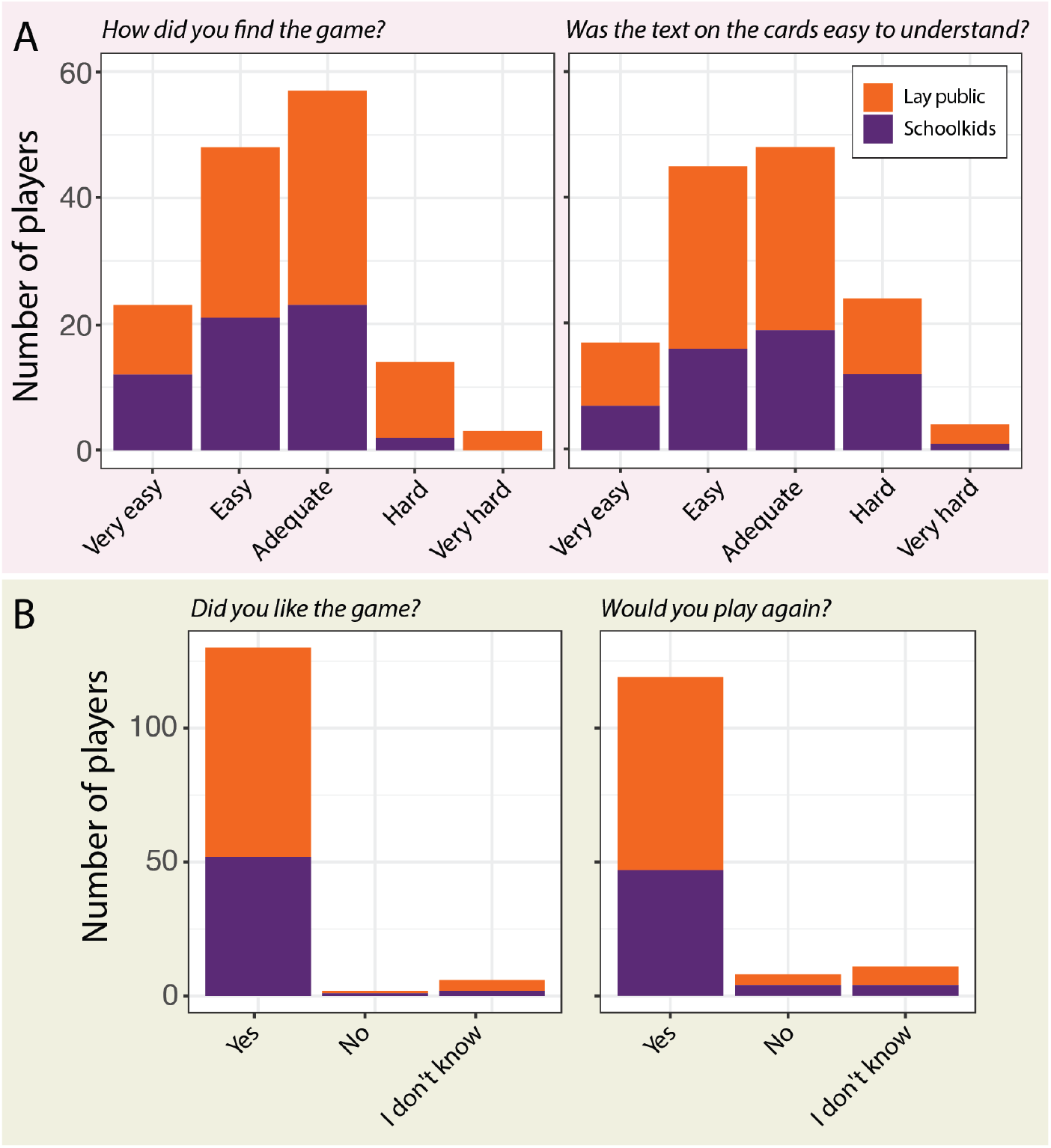
Appreciation of Bacttle by first-time players. A. Feedback on the difficulty of the game and the comprehensibility of the text on the cards. B. General appreciation of the game based on likability and desirability to play again.

## 5. Discussion

Bacttle increases microbiology knowledge on players as young as 5 years old. Based on the positive results observed among players 8-13 years old that played without support from adults, we encourage the implementation of the game as an educational tool in such informal environments as home where children can play on their own. Assuming that the higher improvement score observed in lay public compared to schoolkids is mostly due to the fact that children were playing alongside their family, these results reinforce the potential that this game can have in an educational set up like schools or as a family game at home, where adult support can engage young players into the science of the game more deeply.

In this work, we tested the declarative knowledge acquired immediately after a brief exposure to the game. However, players may forget the facts they could have learnt during the game after a while. Knowledge retention(12) was not specifically tested in this study due to restrictions in collecting personal information about the players, since most of them were minors. Nonetheless, it remains a valuable research question worth considering in the future, if possible.

There are multiple limitations to interpreting research that is based on the answers of people to a survey. The most relevant in our case is the social desirability bias(14), where players will answer what they think is expected from them, rather than what they really think. This bias would have affected the questions regarding basic microbiology knowledge (questions B and C, Table 1), where they were asked if they were familiar with two microbiology concepts. In addition to that, the fact that players were categorized in these groups (‘no knowledge’, ‘basic knowledge’, and ‘advanced knowledge’) based on only two questions limits the accuracy of their actual level of pre-existing microbiology knowledge.

The fact that most players were children adds other biases, for example, their limited verbal and cognitive skills, which could affect their capability to adequately understand and respond to the survey questions. We addressed this bias by preparing short questions and simple vocabulary, but still needed to expose some scientific concepts. Another bias is the reliability of the answers. We could observe this explicitly when a 9-year-old listed an age of 64. Surveys with such obvious erroneous information were discarded, but we may have missed others. One bias that we also observed was parental influence, which we witnessed on a couple of occasions. We tried to address this by separating responses into two groups: schoolkids, where kids played without adult supervision, and lay public, where we observed such possible bias. By comparing our data to a random simulation of replies, we controlled to some extent poor reliability, parental influence bias and other random errors. All results and controls clearly pointed to a positive learning effect of the game on microbiology concepts.

The revised version of the game is illustrated by Philippe Piccardi, and includes the improvements proposed by the players (e.g. simplified text on the cards). The game includes a minimalistic rules manual with mostly diagrams and little text, to ease understanding for young players. Moreover, the way the game has been designed allows for addition of new environmental challenges and traits in future editions. Lastly, Bacttle will be available as open-source material, printable at home, but we are also looking for professional producers of the game. To be updated on the news or latest versions of Bacttle, please refer to www.bacttle.com.

## 6. Acknowledgments

The authors acknowledge support from the Swiss National Science Foundation Sinergia program (contract number CRSII5_189919/1), and from the National Centre in Competence Research (NCCR) in Microbiomes (grant number 180575).

We further acknowledge Philippe Piccardi for his work on the design and illustrations for the game, as well as his invaluable feedback, Valentina Benigno for helping us collect the data during Mystères de l’UNIL, Nathan Schaeffer for his contributions to the statistical analysis and simulation, Yannick Rochat (Game lab Lausanne) for his time and advice, Robin Tecon for all his support through the NCCR Microbiomes and, last but not least, all our colleagues who helped us testing the game infinite times to make it better.

## References

1. Abt CC. Serious Games [Internet]. Viking Press; 1970. (Viking compass book). Available from: https://books.google.ch/books?id=5z-QAAAAIAAJ

2. Michael DR, Chen S. Serious Games: Games that Educate, Train and Inform [Internet]. Thomson Course Technology; 2006. Available from: https://books.google.ch/books?id=49kTAQAAIAAJ

3. Zyda M. From visual simulation to virtual reality to games. Computer. 2005 Sep;38(9):25–32.

4. Riopel M, Nenciovici L, Potvin P, Chastenay P, Charland P, Sarrasin JB, et al. Impact of serious games on science learning achievement compared with more conventional instruction: an overview and a meta-analysis. Stud Sci Educ. 2019 Jul 3;55(2):169–214.

5. Gelbart H, Brill G, Yarden A. The Impact of a Web-Based Research Simulation in Bioinformatics on Students’ Understanding of Genetics. Res Sci Educ. 2009 Nov;39(5):725–51.

6. Kiboss JK, Ndirangu M, Wekesa EW. Effectiveness of a Computer-Mediated Simulations Program in School Biology on Pupils’ Learning Outcomes in Cell Theory. J Sci Educ Technol. 2004;13(2):207–13.

7. Rutten N, van Joolingen WR, van der Veen JT. The learning effects of computer simulations in science education. Comput Educ. 2012 Jan;58(1):136–53.

8. Coil DA, Ettinger CL, Eisen JA. Gut Check: The evolution of an educational board game. PLOS Biol. 2017 Apr 28;15(4):e2001984.

9. de Almeida LG, Pasternak Taschner N, Lellis-Santos C. Outbreak! an Online Board Game That Fosters Collaborative Learning of Viral Diseases. J Microbiol Biol Educ. 2021 Apr 30;22(1):ev22.1.2539.

10. McOwat K, Stanley-Wall NR. Biofilm Building: A Simple Board Game to Reinforce Knowledge of Biofilm Formation. J Microbiol Biol Educ. 2018 Mar;19(1):19.1.70.

11. Miller LM, Moreno J, Estrera V, Lane D. Efficacy of MedMyst: an Internet Teaching Tool for Middle School Microbiology. Microbiol Educ. 2004 May;5(1):13–20.

12. Wouters P, van Nimwegen C, van Oostendorp H, van der Spek ED. A meta-analysis of the cognitive and motivational effects of serious games. J Educ Psychol. 2013 May;105(2):249–65.

13. Basler M, Pilhofer M, Henderson GP, Jensen GJ, Mekalanos JJ. Type VI secretion requires a dynamic contractile phage tail-like structure. Nature. 2012 Mar;483(7388):182–6.

14. Fisher RJ. Social Desirability Bias and the Validity of Indirect Questioning. J Consum Res. 1993;20(2):303–15.

